# A Comparison of Techniques to Determine Active Motor Threshold for Transcranial Magnetic Stimulation Research

**DOI:** 10.1101/2024.04.19.590327

**Authors:** Jonathan P. Beausejour, Jay Rusch, Kevan S. Knowles, Jason I. Pagan, Meredith Chaput, Grant E. Norte, Jason M. DeFreitas, Matt S. Stock

## Abstract

The determination of active motor threshold (AMT) is a critical step in transcranial magnetic stimulation (TMS) research protocols involving voluntary muscle contractions. As AMT is frequently determined using an *absolute* electromyographic (EMG) threshold (e.g., 200µV peak-to-peak amplitude), wide variation in EMG recordings across participants has given reason to consider a *relative* threshold (e.g., = 2× background EMG) for AMT determination. However, these approaches have not been systemically compared. **PURPOSE:** We sought to compare the AMT estimations derived from absolute and relative criteria commonly used to determine AMT in the quadriceps muscles, and assess the test-retest reliability of each approach (absolute = 200µV vs. relative = 2× background EMG). **METHODS:** Eighteen young adults (9 males and 9 females; mean ± SD age = 23 ± 2 years) visited the research laboratory on two occasions. All testing was conducted on the dominant limb. During each laboratory visit, maximal voluntary isometric contraction (MVIC) quadriceps torque was measured, with all subsequent TMS procedures conducted as participants maintained 10% of MVIC torque. AMT was derived from each criterion using motor evoked potentials recorded from the vastus lateralis (VL) and defined as the lowest stimulator output (SO%) needed to meet the specified criteria within ≥ 5/10 pulses. The order of criteria (i.e., absolute vs. relative) was randomized during the first laboratory visit, and counterbalanced during the second visit. A paired samples *t*-test, 95% confidence intervals and the effect size were used to compare mean differences in AMT estimations obtained from each criterion during the second laboratory visit. Paired samples *t*-tests, intraclass correlation coefficients (ICC_2,1_), standard errors of measurement (SEMs), and the minimal difference (MD) scores were calculated to assess test-retest reliability of each AMT criterion. **RESULTS:** Differences between the AMT criteria were small and not statistically significant (absolute criterion mean = 48.9%, relative criterion mean = 47.4%; *p* = .309, Cohen’s *d* = 0.247). The absolute criterion demonstrated moderate to excellent reliability (ICC_2,1_= .866 [0.648 – 0.950], SEM = 7.9%, MD = 10.4%), but higher AMTs were observed in the second visit compared to the first (*p =* 0.043). The relative criteria demonstrated good-to-excellent test-retest reliability (ICC_2,1_= .894 [0.746 – 0.959], SEM = 6.9%, MD = 8.9%) and AMTs were not different between visits (*p =* 0.420).

**CONCLUSION:** Quantifying AMT with an absolute voltage threshold of 200µV peak-to-peak amplitude and a relative voltage threshold 2× background EMG resulted in similar estimations within a single testing session. However, the relative voltage criterion demonstrated superior test-retest reliability. TMS researchers aiming to track corticospinal characteristics across visits should consider implementing relative criterion approaches during their AMT determination protocol.

## Introduction

Transcranial magnetic stimulation (TMS) is a non-invasive tool used to focally activate cortical regions of the brain.^1^ Pioneered by Barker, TMS procedures are commonly implemented as an investigatory tool to analyze focal neuronal activity and their associated cortical and subcortical pathways.^3^ In the context of exploring cortical pathways directly involved in voluntary movement, TMS stimulation over the primary motor cortex (M1) has provided important insights into the excitatory and inhibitory properties of the corticospinal tracts. While these techniques are becoming more common in neuromuscular research, critical methodological considerations for TMS protocols remain poorly standardized.^4^ Difficulties in reproducing previously published TMS work, especially those involving voluntary contractions of large lower limb musculature, highlight the lack of methodological consensus among TMS investigations.^5^ With this, objective justification for TMS procedures is warranted to facilitate the adoption of more robust, evidence-based TMS protocols in future investigations.

An important methodological step in TMS procedures is standardizing the strength of the TMS stimulation pulse across participants.^6^ This standardization is based on a participant’s measured response threshold (i.e., motor threshold), which is conditionally defined as the lowest stimulator output intensity (SO%) necessary to evoke a reliable response in the target muscle.^1,7^ These responses are quantified through evaluation of series of peak-to-peak motor evoked potential (MEP) amplitudes recorded with surface electromyography.^8,9^ A widely adopted method used to determine motor threshold is the relative frequency procedure.^6^ The relative frequency procedure uses multiple TMS pulse series of 5-10 pulses to systematically identify the motor threshold, which is ultimately defined as the lowest SO% that results in “positive” MEP responses ≥ a pre-determined voltage threshold in at least half of the pulses administered in a thresholding pulse series (i.e., 5 out of 10 “positive” MEP responses).^7^ Although the relative frequency procedure is commonly adopted due to its intuitive implementation, other thresholding techniques using mathematical convergence models are also considered to reduce the number of pulses needed to determine motor thresholds.^6,10–12^ Once the motor threshold is determined, subsequent stimulation intensities are normalized to this value to measure key cortical and corticospinal characteristics such as excitability, facilitation, and inhibition.^13,14^

To our knowledge, Rossini et al. were among the first to propose a standardize paradigm to estimate motor thresholds in resting and active muscles.^7,15^ Within their active motor thresholding criteria recommendations, they proposed to define a “positive” motor responses to stimulation as absolute MEP values of ≥ 200µV (active motor threshold [AMT]). Although this absolute voltage threshold lacks empirical justification, it is still widely adopted across TMS studies involving quadriceps muscles.^6^ Concerns with using an absolute voltage value to qualify reliable MEP responses in active muscles are well founded in the neuromuscular literature, as there are several known contributories that influence voluntary sEMG signals aside from peripheral muscular contraction.^16^ Specifically, variations in sEMG interelectrode distance and electrode size, muscle size, skin conductivity, and subcutaneous fat thickness may further confound AMT assessments and question the conventional use of absolute threshold criteria. Difficulties in standardizing TMS procedures across published investigations are also confounded by the use of different EMG systems to quantify MEPs.^17^

To improve standardization procedures, quantification of “positive” MEP voltage thresholds as a relative percentage of participants’ voluntary activation at a given force intensity has been proposed. Early adoption of this approach was reported in an investigation by Damron et al., as they reported to use a relative threshold value criterion of 2× background EMG noise at 15% target force intensity to qualify “positive” MEP responses in the flexor carpi radialis muscle^18^. Even though both relative and absolute criteria have been used in the quadriceps,^14,19–21^ no studies have directly compared these approaches to assess whether quantifying MEP responses to participant-specific background EMG produces more robust and precise AMT estimations compared to using an absolute voltage value. Our purpose was to compare AMT estimations derived from relative and absolute criteria (AIM 1), and assess the test-retest reliability within each approach (AIM 2). Given the exploratory nature of this investigation, we hypothesized that AMT estimations would be lower for the relative criterion compared to the absolute criterion. We also anticipated that both criteria would exhibit at least “moderate” reliability.

## Methods

### Experimental Design

A repeated measures design was implemented to compare acquired AMT estimations between the two determination criteria (AIM 1) and assess the test-retest reliability of each criterion (AIM 2). Study participants were required to visit the laboratory on two separate occasions to complete data collection.

During each visit, AMT of the participants’ dominant vastus lateralis (VL) was determined using the absolute (200 µV) and relative (2× background EMG) approaches consecutively. The order of AMT determination criteria used during the first visit was randomized, and subsequently counterbalanced for the second visit. Data collection for each participant was completed at the same time of day (± 1 hour) and laboratory visits were separated by at least 48 hours, but no longer than two weeks. Each laboratory visit lasted between 1.5 to 2 hours. Given the possibility of induced muscular fatigue, three-minute rest periods were administered after maximal torque testing and TMS hotspot testing, one-minute rest periods were administered between each series of TMS pulses, and a 10-minute rest period was given between each AMT determination criterion. All participants were asked to refrain from moderate-to-strenuous exercise and alcohol consumption for at least 24 hours, and caffeine consumption at least 4 hours, prior to data collection.

### Participants

Eighteen healthy, young adults (nine males, nine females; mean ± SD age = 22 ± 2 years, height = 170.7 ± 10.4 cm, mass = 72.4 ± 18.1 kg, body mass index [BMI] = 24.6 ± 4.2 kg/m^2^) completed the study. Inclusion criteria included healthy adults between 18 and 30 years of age and BMI between 18.5-35.0 kg/m^2^. Exclusionary criteria included a history of concussions, stroke, heart attacks, cancer, presence of major neuromuscular/metabolic diseases, family history of epilepsy and/or syncope, and lower-body musculoskeletal injury and/or surgery within the previous six months. Participants were also excluded if they were taking prescription medication known to affect physiological properties of cortical excitability (e.g., alpha, beta, and/or calcium channel blockers, sedatives, psychostimulants).^22^ All participants signed informed consent documents, completed the physical activity readiness questionnaire, and underwent comprehensive TMS screening prior to data collection.^22^ The study was approved by the University Institutional Review Board (#4836).

### sEMG Signals

sEMG signals were recorded from the participant’s dominant VL muscle (based on their preferred limb to kick a ball) with a bipolar surface EMG sensor (Trigno™ EMG, Delsys, Inc., Natick, MA, USA; interelectrode distance = 10mm, bandpass filter: 20-450 Hz). The EMG sensor was placed at 2/3 the distal distance between the greater trochanter and the superolateral border of the patella.^23^ During sensor placement, participants were asked to lightly contract their quadriceps muscles to precisely identify the VL muscle belly at the 2/3 distance. The skin over the belly was shaved and cleansed with rubbing alcohol, and hypoallergenic tape was used to remove any excess hair and dead skin cells. A permanent marker was used to denote the location of the sEMG sensor for the subsequent laboratory visit. sEMG signals were sampled at 2 kHz, and a signal to noise ratio > 3.0 and line interference < 1.0 were confirmed during a light, voluntary contraction of the quadriceps muscles to ensure high-quality signal acquisition (EMGworks acquisition software, version 4.7.6, Delsys, Inc., Natick, MA, USA).

### Isometric Peak Torque

All maximal/submaximal voluntary isometric contractions were completed on the participants’ dominant quadriceps muscles with a Biodex System 4 isokinetic dynamometer (Biodex Medical Systems, Inc., Shirley, NY, USA). Participants were secured to the dynamometer via straps placed across the hips and chest in a “X” pattern. The dominant lower leg was strapped in place so that their knee joint was in line with the dynamometer axis attachment. Knee and hip joint angles were set 110° and 95°, respectively (180° = full extension). Following a brief warm-up, participants performed three, five-second maximal voluntary isometric contractions (MVIC). At least two minutes of rest was provided between MVIC trials, and the highest peak torque value was used for subsequent quantification of 10% submaximal torque intensity during the active TMS protocol. Participants were provided visual feedback of their torque performance during all maximal and submaximal contractions.

### Single Pulse TMS

Single-pulse TMS was administered over the participants’ primary motor cortex (M1) with a Magstim®200² stimulator device and corresponding 110 mm double cone coil (The Magstim Company Ltd, Whitland, UK). An anterior-posterior orientation was utilized throughout the protocol. Upon completion of the initial MVIC trials, participants were given a disposable Lycra swim cap to place over their head. The vertex for each participant was demarcated (via black permanent marker) by identifying the intersection point between lines spanning from the occipital protuberance to the nasal, and from right tragus (ear) to left tragus. Next, a 1 × 1-centimeter grid were marked on the swim cap lateral to the vertex, and contralateral to the side of the participants’ dominant limb. Hotspot determination started two centimeters lateral to the vertex (contralateral to the side of the participants’ dominant limb), and progressed by methodically testing each adjacent grid point (in a randomized order) with five TMS pulses. Participants performed isometric knee extension contractions at 10% MVIC during all hotspot stimulation trials, and each hotspot trial were completed at 40% SO. The hotspot location was identified as the grid point that elicited the highest mean MEP amplitude response in the VL and marked with a red permanent marker. When necessary, new points were added to the grid to ensure all viable locations were tested. To ensure marker placement consistency across visits, participants wore the same swim cap for their second laboratory visit and fitted to ensure proper alignment of their predetermined vertex and grid locations. Hotspot determination was re-assessed in the second laboratory visit. Location of the windings that corresponded to the determined M1 hotspot was also marked on the Lycra cap to ensure consistent coil placement throughout testing. The same investigator (J.P.B.) held the TMS coil across all testing visits.

After hotspot determination, AMT testing was conducted at the hotspot location using the absolute (200 µV) and relative (2× background EMG) approaches consecutively. For each approach, AMT was defined as the lowest stimulator output (SO%) that met the respective MEP voltage criterion in at least 5 out of 10 pulses. A custom-LabVIEW program (NI LabVIEW 17.0, National Instruments, Austin, TX, USA) was used to provide visual torque tracings for participants to perform active contractions at 10% MVIC force, and real time quantification of TMS-derived VL MEP responses during each determination pulse series (10 pulses per series). For each pulse series, the visual torque tracing was programmed to provide a trapezoidal pattern corresponding to 10 instances in which participants were to contract their dominant limb quadriceps muscles at 10% MVIC force. The duration of each contraction was 4 seconds, with 5-second rest intervals and 0.5 seconds ramping up to and down from the 10% MVIC force, respectfully. Background EMG was quantified as the highest peak-to-peak amplitude observed 25 milliseconds (ms) before the deliverance of the TMS pulse. To further standardize the TMS protocol across participants and track number of pulse series needed to determine AMT, TMS stimulation began at 40% SO for each criterion. Following each pulse series, the TMS SO% was systematically adjusted based on the number of “positive” MEP responses’ absolute difference from 5. For example, if a pulse series resulted in 1/10 MEPs that met the respective criterion, the SO% intensity would have increased by 4 for the next 10-pulse series. Conversely, if a pulse series resulted in 7/10 MEPs that met the respective criterion, the SO% intensity would have reduced by two for the next 10-pulse series. Inter-pulse interval duration during AMT determination trails were between 7 to 10 seconds, as priority was given for participants’ precise torque tracing at 10% MVIC force, during each 4-second contraction instance, before deliverance of the pulse. Following data collection, participants performed three more MVIC trials to assess the level of performance fatigue induced by the TMS protocol.

### Statistical Analysis

To examine differences in AMT between the criteria, a paired samples *t*-test (2-tailed) and 95% confidence intervals (CIs) were used for data obtained during the second visit. Pre to post changes in MVIC peak torque during the second visit were also assessed using a paired samples *t*-test. An alpha level of 0.05 was utilized for all analyses. Cohen’s *d* values of 0.2, 0.5, and 0.8 were used to denote either a small, moderate, and large difference, respectfully.^24^ A Bland-Altman plot was used to identify 95% limits of agreements (LOA) between the AMT criteria for the second laboratory visit.^25^

Test-retest reliability for each AMT criterion was quantified with a 2-way random-effects intraclass correlation coefficient model (ICC). This model was used, specifically, to report generalizable reliability metrics for future TMS research.^26^ ICCs were evaluated based on the 95% confident intervals of the ICC point estimation, in which ICC values < 0.50 indicated “poor” reliability, ICCs between 0.50 - 0.75 were indicative of “moderate” reliability, ICCs between 0.75 - 0.90 were indicative of “good” reliability, and ICCs > 0.90 were indicative of excellent reliability.^27^ The mean square error was used to determine the Standard error of the measurement ([SEM] expressed in absolute units and percentage of the grand mean), and the minimal difference value needed to be considered real (MD). In addition, paired samples *t*-tests (2-tailed) were used to examine mean differences in AMT within each criterion, along with 95% confidence intervals for the mean differences and Cohen’s *d* effect sizes. Assumptions of normality were verified via Shapiro-Wilks tests prior to statistical interpretation.^28^ JASP statistical software (JASP 16.0, University of Amsterdam, Amsterdam, NL) was used to conduct all paired samples *t*-tests, and ICCs. SEM and MD values were calculated based on the formulas outlined in the Weir (2005) report.

## Results

### Between criteria comparison (AIM 1)

**Table 1** presents AMT estimations for each participant during the second visit, as well as descriptive data for the MEP peak-to-peak amplitudes recorded at the AMT intensity level for each criterion. There was no difference in the AMT observed between the criteria (*p* = 0.309) and the effect size was small (*d* = 0.247). The mean difference in AMT was 1.44 units (95% CI: [-1.461 – 4.350]), with slightly higher AMT observed in the absolute criterion compared to the relative criterion (95% LOA = [- 10.01 – 12.90]) (**Figures 1 and 2**). Comparison of the number of pulse series needed to acquire AMT did not differ between the criterion (200µV mean ± SD = 5.0 ± 2.5, 2× background EMG = 5.3 ± 2.0; *p =* 0.588; *d =* 0.130).

**Table 1.** Visit 2 AMT descriptive statistics, including the acquired AMT estimations, the mean ± SD of recorded MEP responses at the determined threshold series (10 pulses), and the number of pulse series needed for AMT determination for each criterion.

| Participant | AMT<br>(200μV) | AMT<br>(2× Background) | MEP Response<br>at AMT<br>(200μV) | MEP Response at<br>AMT<br>(2× Background) | Number of AMT<br>Hunting Trails<br>(200μV) | Number of AMT<br>Hunting Trails<br>(2× Background) |
| --- | --- | --- | --- | --- | --- | --- |
| AMT 001 | 55 | 60 | 224.1 ± 54.7 | 258.9 ± 60.0 | 7 | 7 |
| AMT 002 | 40 | 39 | 212.3 ± 43.6 | 189.5 ± 31.9 | 2 | 3 |
| AMT 003 | 44 | 35 | 233.8 ± 69.4 | 83.6 ± 28.7 | 3 | 6 |
| AMT 004 | 50 | 56 | 274.2 ± 83.7 | 461.8 ± 209.0 | 5 | 9 |
| AMT 005 | 38 | 48 | 334.7 ± 81.9 | 601.8 ± 252.1 | 3 | 4 |
| AMT 006 | 73 | 65 | 228.7 ± 69.7 | 98.6 ± 34.6 | 11 | 7 |
| AMT 007 | 44 | 45 | 256.2 ± 68.5 | 200.6 ± 68.4 | 5 | 4 |
| AMT 009 | 40 | 42 | 234.5 ± 58.2 | 297.0 ± 65.6 | 2 | 3 |
| AMT 011 | 56 | 45 | 215.5 ± 30.7 | 76.33 ± 17.8 | 6 | 4 |
| AMT 012 | 49 | 47 | 259.6 ± 73.0 | 237.1 ± 70.8 | 5 | 4 |
| AMT 013 | 40 | 35 | 215.1 ± 53.8 | 150.0 ± 46.0 | 2 | 5 |
| AMT 014 | 75 | 66 | 209.4 ± 55.3 | 85.6 ± 38.1 | 9 | 10 |
| AMT 015 | 57 | 51 | 299.6 ± 155.1 | 72.0 ± 30.8 | 7 | 5 |
| AMT 016 | 37 | 37 | 230.1 ± 33.2 | 224.6 ± 50.4 | 5 | 6 |
| AMT 017 | 40 | 41 | 253.8 ± 92.0 | 244.2 ± 71.6 | 3 | 3 |
| AMT 018 | 50 | 52 | 248.2 ± 70.7 | 239.9 ± 95.4 | 5 | 6 |
| AMT 019 | 42 | 37 | 209.1 ± 38.7 | 75.5 ± 21.6 | 3 | 5 |
| AMT 020 | 50 | 53 | 202.3 ± 57.6 | 270.1 ± 166.2 | 7 | 4 |

**Figure 1.**
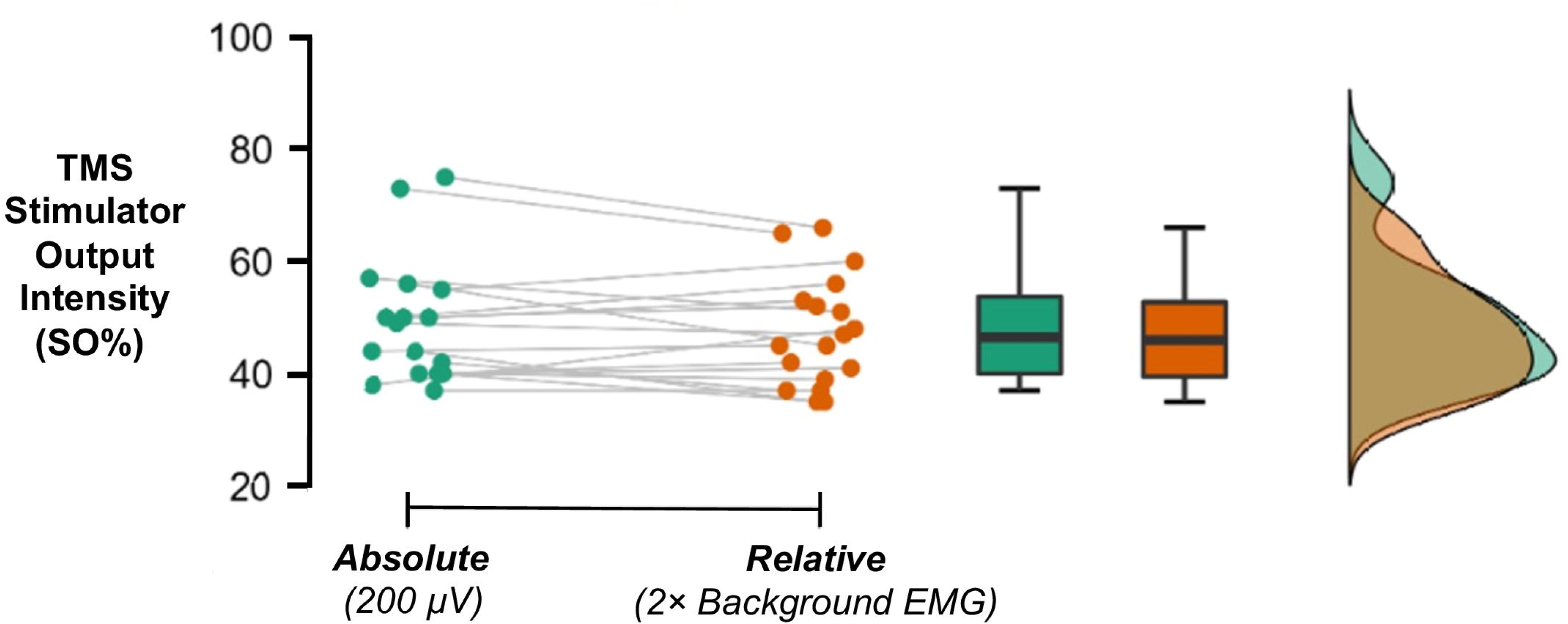
Raincloud plot of AMT estimations acquired for each criterion. Each circle and line correspond to data for an individual participant (n = 18). Data distribution plots (box-and-whisker and histogram) display the normally distributed data for each AMT criterion. The results indicated that differences between AMT were small and not statistically significant.

**Figure 2.**
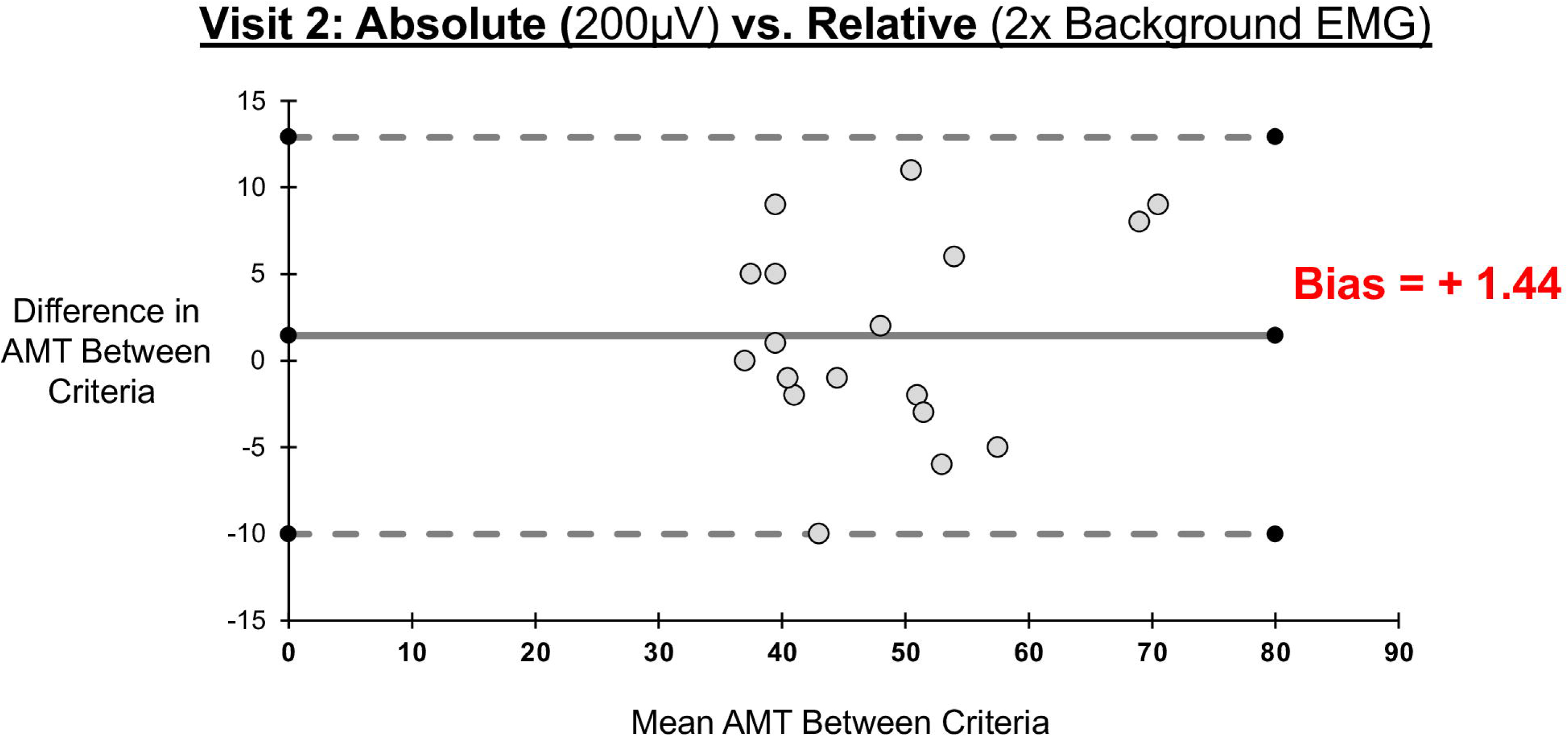
A Bland Altman plot displaying agreement between the AMT criteria during the second laboratory visit. The x-axis represents the mean AMT acquired from each criterion. The y-axis represents the mean difference between the criterion during the second laboratory visit, with the bias denoting the mean difference of the absolute criterion compared to the relative criterion. Dotted lines: 95% limits of agreement (LOA) = [-10.09 – 12.90].

### Within criteria reliability (AIM 2)

**Table 2** and presents the reliability statistics for AMT acquired in each criterion. The mean ± SD duration between laboratory visits was 5.5 ± 3.6 days (range: 2 – 14 days). The absolute criterion displayed moderate-to-excellent reliability (ICC = .866 [0.648 – 0.950]) and the relative criteria displayed good-to-excellent reliability (ICC = .894 [0.746 – 0.959]). The relative criteria exhibited slightly lower SEM values compared to the absolute criterion (absolute = 3.74, 7.92%; relative = 3.23, 6.94%). Mean AMT estimations were higher for visit 2 than visit 1 when using the absolute criterion (*p =* 0.043; mean difference = 2.72; 95% CI: [0.09 – 5.35]; *d =* 0.515). AMT for the relative criterion was not different between visits (*p* = 0.420, mean difference = 0.889; 95% CI: [-1.380 – 3.157]; *d* = 0.195) (**Figure 3)**.

**Table 2.**
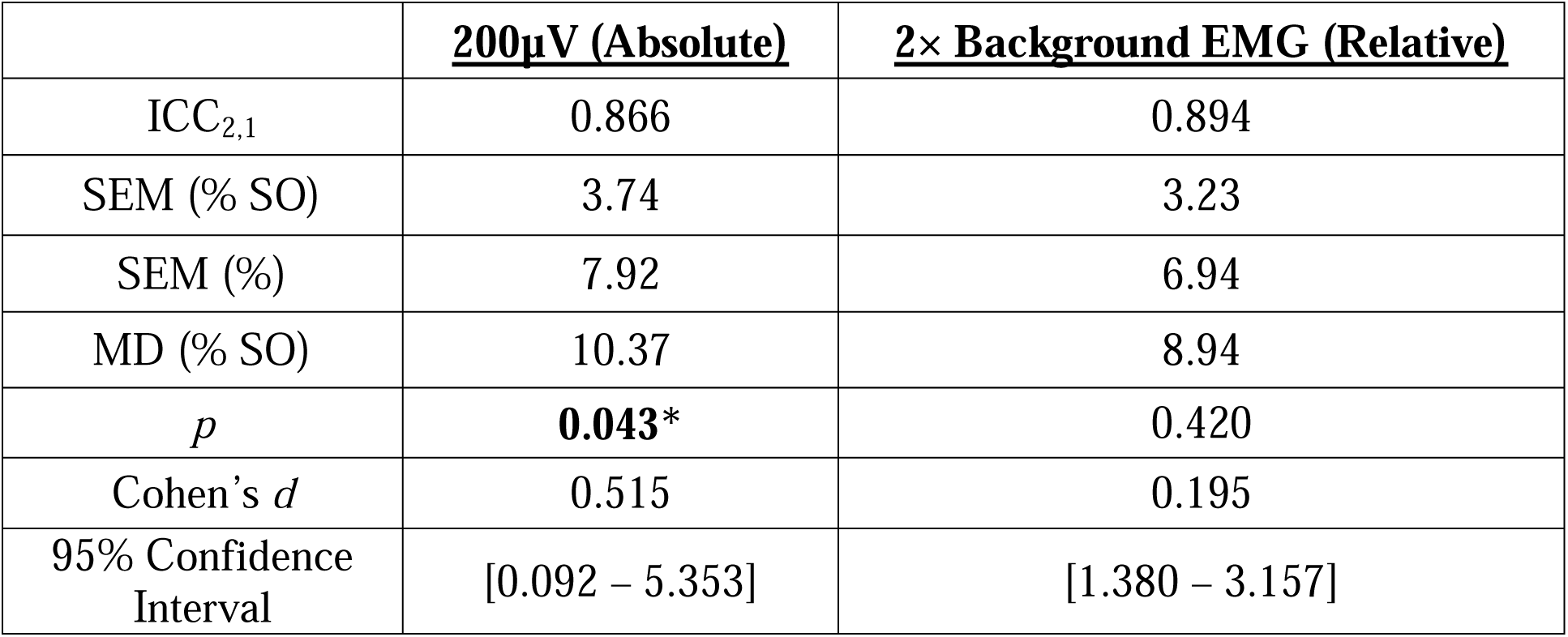
ICC and paired samples *t*-test results. Intraclass Correlation Coefficients (ICCs) and associated 95% confident intervals, Standard Error of Measurements (SEMs), and Minimal Difference Change scores (MD) for AMT estimations acquired via the absolute (200µV) and relative (2× background EMG) criteria. P values, 95% confidence intervals for mean difference, and Cohen’s d effect sizes for paired samples *t*-tests conducted for each criterion are also displayed. \**p* < .05.

|  | **<u>200μV (Absolute)</u>** | **<u>2× Background EMG (Relative)</u>** |
| --- | --- | --- |
| ICC<sub>2,1</sub> | 0.866 | 0.894 |
| SEM (% SO) | 3.74 | 3.23 |
| SEM (%) | 7.92 | 6.94 |
| MD (% SO) | 10.37 | 8.94 |
| *p* | **0.043\*** | 0.420 |
| Cohen’s *d* | 0.515 | 0.195 |
| 95% Confidence Interval | [0.092 – 5.353] | [1.380 – 3.157] |

**Figure 3.**
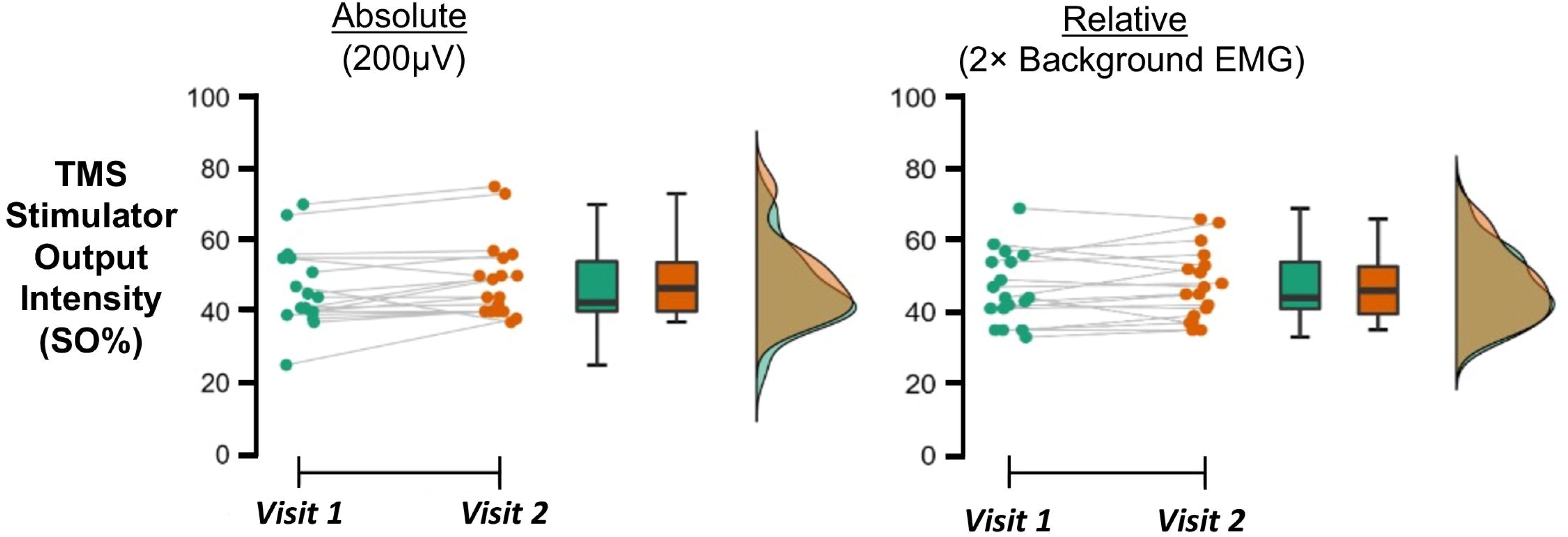
Raincloud plot of AMT estimations acquired in both laboratory visits for each criterion. This plot presents repeated measures data for each participant, with a trendline reflecting the individual change in acquired values.

### Pre-Post MVIC Peak Torque

Figure 4 presents individual participant MVIC peak torque performance before and after the AMT protocol for visit 2. Results from the paired samples *t*-test indicated that quadriceps MVIC peak torque did not change following the TMS protocol (pretest MVIC mean ± SD = 220.1 ± 72.4 Nm, posttest MVIC = 221.2 ± 75.3 Nm; *p =* 0.761), and the effect size was small (*d =* 0.073).

**Figure 4.**
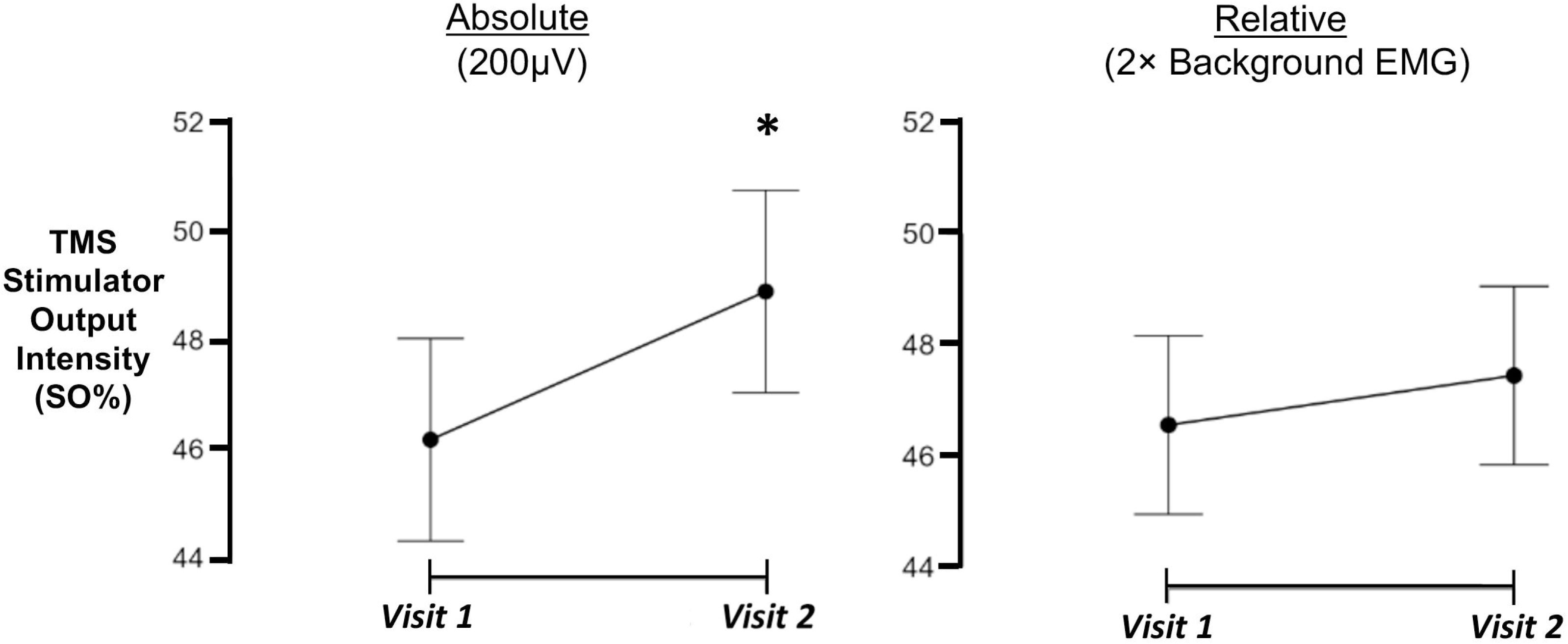
Descriptive plots displaying the mean ± standard error for each criterion. A trendline reflects the change in AMT mean values between visits. \**p* < .05.

**Figure 5.**
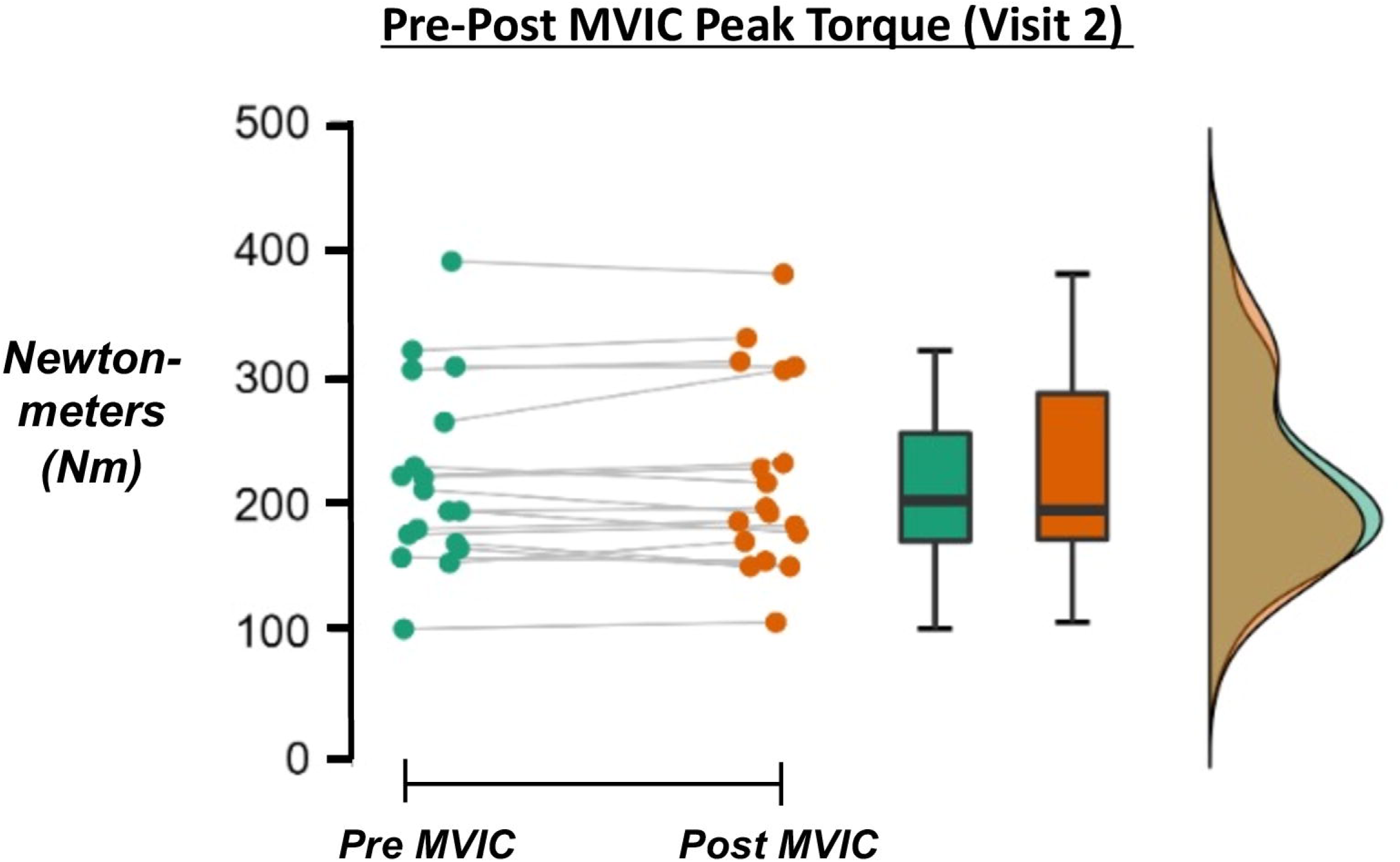
Raincloud plot of pre-post MVIC performance (expressed in newton-meters [Nm]) for the second laboratory visit. This plot displays pre-post MVIC performance for each participant, with a trendline reflecting individual change before and after the TMS protocol.

## Discussion

AMT determination is a critical methodological step in TMS research protocols that involves active contractions. We sought to compare two voltage-based criteria commonly used to determine AMT in the quadriceps, and assess the test-retest reliability of each approach. Although our findings show that similar AMTs were found for both the absolute and relative criteria, reliability metrics differed between the approaches. MVIC peak torque was also consistent following the TMS protocol, thus indicating that the protocol was not fatiguing. Below we discuss our interpretation, limitations, and noteworthy perspectives.

Concerns in adopting absolute voltage criteria for AMT estimations are rooted in the wide variation of sEMG-recorded MEP responses between individuals.^16,29^ This variation is realized through individual differences in cortical conduction mechanisms that contribute to the desynchronized excitation of target muscle(s) from TMS pulses.^29^ Characteristics of this desynchronized excitation may be further altered when active contractions are introduced, as sEMG signals from voluntary contractions are representative of multiple motor unit properties that may vary across participants.^16^ Furthermore, factors like thickness of subcutaneous tissue and the variable distribution of muscle fibers and fiber conduction velocities influence sEMG signal acquisition and cannot be controlled between participants.^16^ However, our results revealed no difference in AMT between the absolute and relative criteria. Rationale for this finding may include the voltage sufficiency of the specific absolute voltage criterion used in this study. As the relative approach of 2× background EMG may be considered to confidently discern MEPs from background activation across participants, an absolute voltage threshold of 200uV may similarly account for the inter-participant variability of MEPs in a single testing session. This is notable, as our investigation is the first to compare these criteria while also controlling for other procedural components included in TMS protocols (e.g., TMS stimulator devices, sEMG electrode placement, and sEMG acquisition systems). Deviations in such components may yield different results than presently observed, as the acquisition of sEMG-recorded MEPs with differing laboratory techniques further complicates AMT estimations across TMS investigations.

Previous reliability studies in healthy participants have reported quadriceps AMT reliability to range between moderate to excellent, despite the different criterion used during AMT determination.^14,20,30^ In accordance with prior studies, the relative approach in our investigation exhibited good-to-excellent reliability, and the absolute approach demonstrated moderate-to-excellent reliability. While mean AMT estimations across visits shifted in a similar direction for each criterion, our results indicated an overestimation of AMT across visits for the absolute approach. As such, precise estimations of AMT may be more effectively accomplished with a relative voltage criterion compared to absolute voltage approaches, as relative approaches may provide more appropriate voltage parameters to qualify MEPs between participants. Considering stimulation intensities employed to explore key cortical and corticospinal often range from 50% to 200% AMT, precise AMT estimations across visits are vital to ensure appropriate testing stimulations in TMS investigations.

With the predominant use of absolute voltage criteria across TMS studies, our results also contribute to recent investigations that report reliable tracking of TMS variables with relative criterion approaches. To our knowledge, only one investigation reported good reliability (ICC = .850) in AMTs acquired from a quadriceps muscle (rectus femoris) using this same relative criterion.^14^ Outside of that investigation, reliability of active TMS variables acquired via relative thresholding criteria in the quadriceps were not explored until the works of Temesi et al. and Shih et al., in which moderate-to-excellent reliability was reported for VL MEPs and specific input/output curve variables during single-pulse stimulation^30,31^. Of note, both the Temesi et al. and Shih et al. investigations used a relative thresholding criterion of “discernable” MEP responses larger than background EMG activation during 20% MVIC, which differed from the relative criterion of 2× background EMG at 10% MVIC employed in our investigation. With multiple absolute and relative AMT voltage criteria reported in TMS studies, future research should aim at investigating how disparities within each voltage criterion influence AMT and other TMS variables across visits.

Considerations regarding AMT reporting and parameters for certain voltage criteria may be evaluated on a case-by-case basis. The SEM and MD scores observed in our study suggest that relative criterion approaches may be more sensitive in detecting AMT changes that can be considered meaningful. Although AMT expresses limited interpretive potential, significant changes in AMT may provide insight into specific neurophysiological adaptions following an intervention.^29^ Selection between absolute and/or relative criteria for AMT may also depend upon specific task-specific conditions. As our investigation assessed AMT estimations during open-chain activation, re-examining these criteria in more functional conditions (i.e., closed-chain) may yield different outcomes. For reference, Young et al. used the TMS Motor Threshold Assessment Tool^12^ to determine AMTs for the vastus medialis and demonstrated poor-to-good reliability during a single let squat condition^30^. It is still unclear whether the voltage approaches used in the present investigation would yield better test-retest reliability during closed-chain activation.

Another important factor is the specific voltage parameters of absolute/relative criteria approaches. Specific voltages for absolute criteria have ranged from 100µV to 200µV for AMT acquired in the quadriceps muscles, and pre-pulse muscle activation levels have been reported anywhere between 5-20% MVIC.^21,32,33^ Opting to select a smaller absolute value (e.g. < 200µV) to qualify VL MEP responses for AMT determination may result in difficulty identifying MEPs from background EMG activation, especially if the pre-pulse activation levels are at 20% MVIC. Alternatively, 200µV may be too high of a voltage threshold for participants whose background EMG activation is substantially smaller. Relative criterion approaches may better accommodate differences in contraction intensities used across TMS studies, and address some of the difficulties in standardizing TMS research participants across visits.

This study has several limitations to consider. Our AMT determination approach is limited to the relative-frequency procedure, and our results may not apply to other AMT thresholding procedures. Our results are also limited to the target muscle (VL), the task employed during TMS (open chain activation on dynamometer), pre-pulse activation level (10% MVIC), and specific parameters used for the absolute and relative voltage criterion (200 µV vs. 2× background EMG). We included only young healthy adults with no neurophysiology and/or musculoskeletal injury pathology, thus limiting our findings to these specific population characteristics. Although permanent markings were placed over the participants’ VL muscle to ensure consistent sEMG sensor placement (as based on the SENIAM guidelines), minor differences in sensor placements may have occurred between visits. We also did not track specific details pertaining to sleep, stress, and arousal levels during the study, yet these variables could influence TMS measures. We also did not attempt to collect specific morphology characteristics of participants’ target muscle (e.g., muscle thickness/cross sectional area, thickness of subcutaneous tissue, segmental body composition). Future TMS studies may consider collecting such data to include as a co-variate, or explore how these data influence MEP responses in the quadriceps. Finally, all of our TMS procedures were performed manually. The use of automated robotic stimulations and/or neuro-navigation software may have alleviated any potential human error during TMS administration.

In conclusion, our results suggest that quantifying AMT with an absolute voltage threshold of 200µV peak-to-peak amplitude and a relative voltage threshold 2× background EMG resulted in similar estimations within a single testing session. However, the relative voltage criterion demonstrated superior test-rest reliability. Based on our findings, we believe that TMS researchers looking to explore corticospinal characteristics across visits should consider implementing relative criterion approaches during their AMT determination protocol. Widespread implementation of relative AMT approaches may minimize variability across laboratories, allow for more consistent communication among TMS researchers, and facilitate the development of consensus guidelines for TMS research.

## Data statement

Data available upon request.

## Compliance with Ethical Standards statement

Study procedures were in compliance with the 1964 Helsinki declaration and its subsequent amendments. Each study participant read, understood, and signed an informed consent document prior to participation.

